# Olfactory Training Enhances Adult Neurogenesis of the Olfactory Epithelium Improving Odor Discriminability of the Olfactory Bulb

**DOI:** 10.64898/2025.12.05.692562

**Authors:** Vittoria Avaro, Elham Taha, Sapir Shapira, Gustav Kühn, Thomas Hummel, Adi Mizrahi, Federico Calegari

## Abstract

Olfaction enables us to detect chemicals in the environment, and its impairment affects one in four people worldwide. As the most common therapy, olfactory training involves repeated exposure to odorants restoring function in 30-50% of patients. However, the mechanisms underlying this therapy, and its partial efficacy, are unknown. By modelling olfactory training in mice, we found that odor exposure triggers quiescent neural stem cells of the olfactory epithelium to re-enter the cell cycle, enhancing neurogenesis and survival of olfactory sensory neurons. Coupling neuronal activity with stem cells proliferation, these effects scaled with odor concentration and were replicated without odorants upon chemogenetic activation of olfactory sensory neurons. Imaging mitral cells in the olfactory bulb revealed that increased neurogenesis in the olfactory epithelium enhanced odor sensitivity and discriminability. Our findings uncover activity-dependent cellular mechanisms linking peripheral nervous system regeneration to central olfactory processing, offering new avenues for treating olfactory dysfunction.

**GRAPHICAL ABSTRACT:** 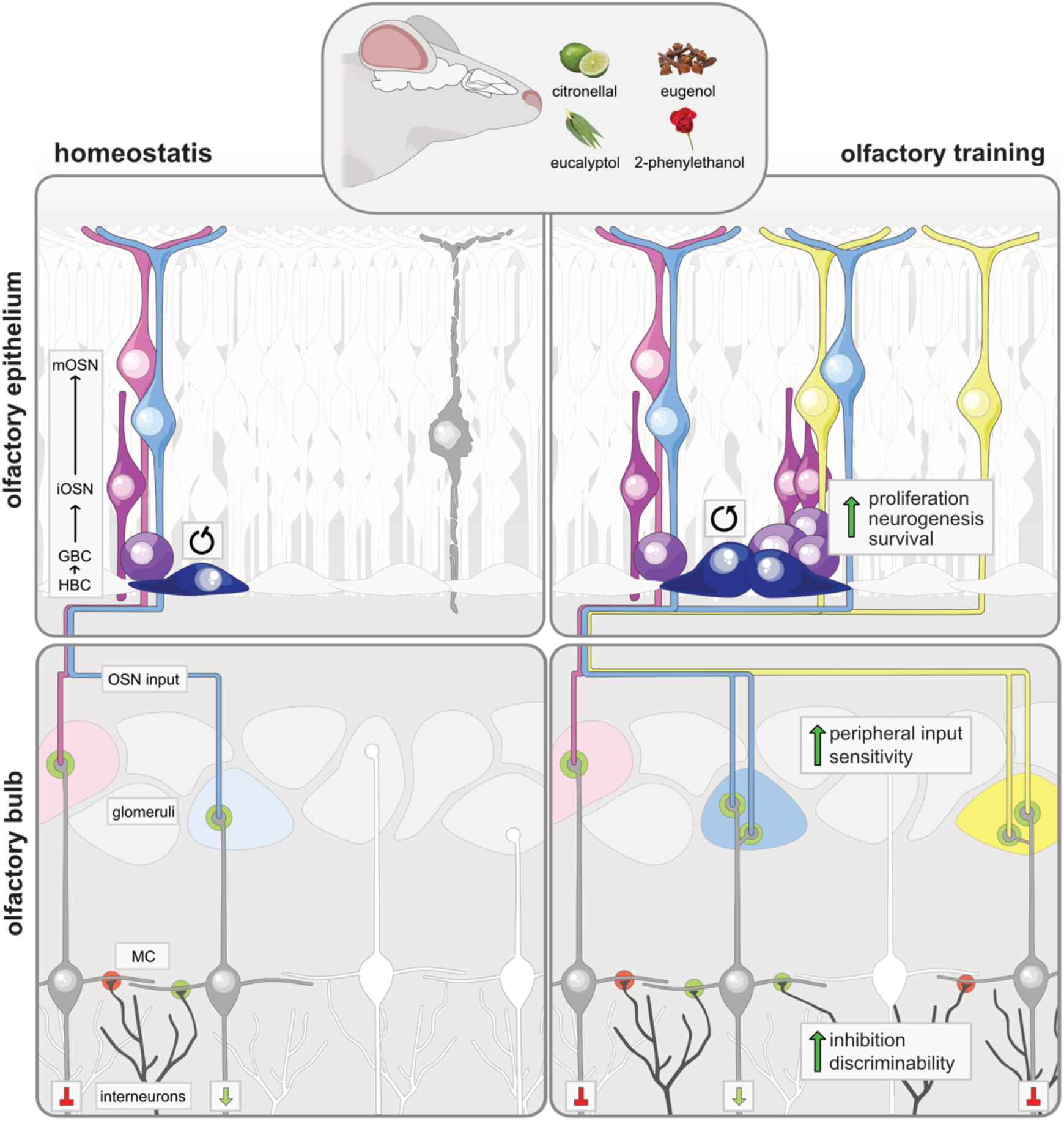

**HIGHLIGHTS:**

- Olfactory training (OT) is a common clinical practice of unknown cellular mechanisms
- OT in mice triggers stem cell activation and neurogenesis in the olfactory epithelium
- Artificial stimulation of olfactory neurons is sufficient to trigger neural stem cell activity
- Neurogenesis in the olfactory epithelium improves odor discrimination by mitral cells

## INTRODUCTION

Olfaction is a highly specialized chemosensory modality, enabling us to perceive thousands of odorants by converting them into biological signals that give rise to our sense of smell^1^. Contrary to common belief, humans possess an exceptional sense of smell that, even if unconsciously, plays critical roles in detecting hazards and regulating appetite, mood and social behaviour^2,3^. Notably, 20-30% of people worldwide experience olfactory dysfunctions^4^ that, while often undiagnosed, underappreciated or untreated, significantly reduce quality of life^5,6^. Linked to depression and early marker of neurodegeneration preceding diagnosis by years of even decades^7^, loss of smell can be caused by congenital defects, chronic rhinosinusitis, head trauma, respiratory infections or aging^8,9^. With an aging global population and recent diseases such as COVID-19, identifying treatments rescuing our sense of smell has become an increasingly important scientific and clinical priority.

Olfactory Training (OT) was characterized in 2009 as the only effective non-pharmacological treatment for most olfactory dysfunctions^10^. OT consists of systematic short-term exposures to a set of odorants, with various training regimens now used worldwide to rescue olfactory function in approximately 30-50% of patients, depending on aetiology^11^. According to the original protocol, patients are instructed to sniff 4 odorants, for 10 seconds each, twice daily for 12 weeks leading to an improvement in olfactory threshold, discrimination and identification^10,11^. Despite its widespread use, the biological mechanisms underlying OT remain unknown. Understanding how olfactory exposure modulates the peripheral and central stations of the olfactory system is critical for developing more effective training regimens for non-responding patients.

Odor detection begins in the Olfactory Epithelium (OE), as part of the peripheral nervous system, where odorants bind to olfactory receptors expressed by Olfactory Sensory Neurons (OSN). Olfactory receptors constitute the largest gene family in our genome, with each neuron expressing only a single receptor gene, resulting in a mosaic-like distribution of OSN subtypes across the OE^12,13^. Tuned to specific molecular features, the combinatorial activation of OSN defines olfactory threshold enabling the detection of thousands of distinct odors. Odor information from the OE is transmitted to the Olfactory Bulb (OB), first station of the central nervous system, where highly plastic processing occurs that dynamically adapts to learning experience and environmental demands^14–16^. Within the OB, OSN terminals synapse onto mitral and tufted cells, which are the principle output neurons of the OB^17^. Mitral Cells (MC) integrate input from OSN with output derived from the local processing of a rich network of inhibitory interneurons^18^ and cortical feedback^19^. Axons of mitral and tufted cells then project to multiple cortical and subcortical areas, where the encoded information supports odor identification and memory recall triggering emotions and guiding behavior^17^. In other words, the OE–OB pathway is thought to provide an initial shaping of odor representations for subsequent computation, integration, and interpretation underlying our sense of smell.

Remarkably, both the peripheral OE and central OB are exceptional niches in that characterized by the continuous integration of adult-born neurons over the course of life^20,21^. The regenerative potential of the OE has long been recognized^22^ and human and rodent neural precursors residing along the basal lamina of the pseudostratified OE have been characterized both at the cellular and molecular level^23,24^. These precursors include Horizontal Basal Cells (HBC) that are predominantly quiescent and cycling Globose Basal Cells (GBC). Under physiological conditions, multipotent HBC function as a long-term reserve of neural stem cells, generating GBC and non-neuronal cell types, such as supporting sustentacular cells^21,24^. Conversely, transit amplifying GBC primarily undergo neurogenic division to generate OSN^21^. While the OE retains the ability to fully regenerate both neuronal and non-neuronal cell types^25,26^, the OB is solely supported by the integration of interneurons generated by neural stem cells residing within the Subventricular Zone (SVZ)^27,28^.

Aiming to support the development of more effective therapies for olfactory dysfunctions, here we investigate the mechanisms underlying OT by establishing an olfactory stimulation protocol in mouse that recapitulates the clinical treatment. Using this model, we assessed the effects of OT on stem cell proliferation, adult neurogenesis and neuronal survival in the peripheral OE while, in parallel, investigating changes in odor processing in the central OB.

## RESULTS

### OT induces quiescent HBC to enter the cell cycle

To investigate the mechanisms underlying OT, we established a paradigm of olfactory stimulation in adult mice that recapitulated the first described human treatment^10^. Specifically, mice were exposed twice a day to four undiluted odorants: citronellal, eugenol, eucalyptol and 2-phenylethanol (cit, eug, euc and 2pe as odors characteristic of lime, clove, eucalyptus and rose, respectively) (Fig. 1A). Two weeks later, the OE was processed for immunohistochemistry and stem cells and their progeny were identified by the expression of their respective markers (Fig. 1B and C).

**Figure 1.**
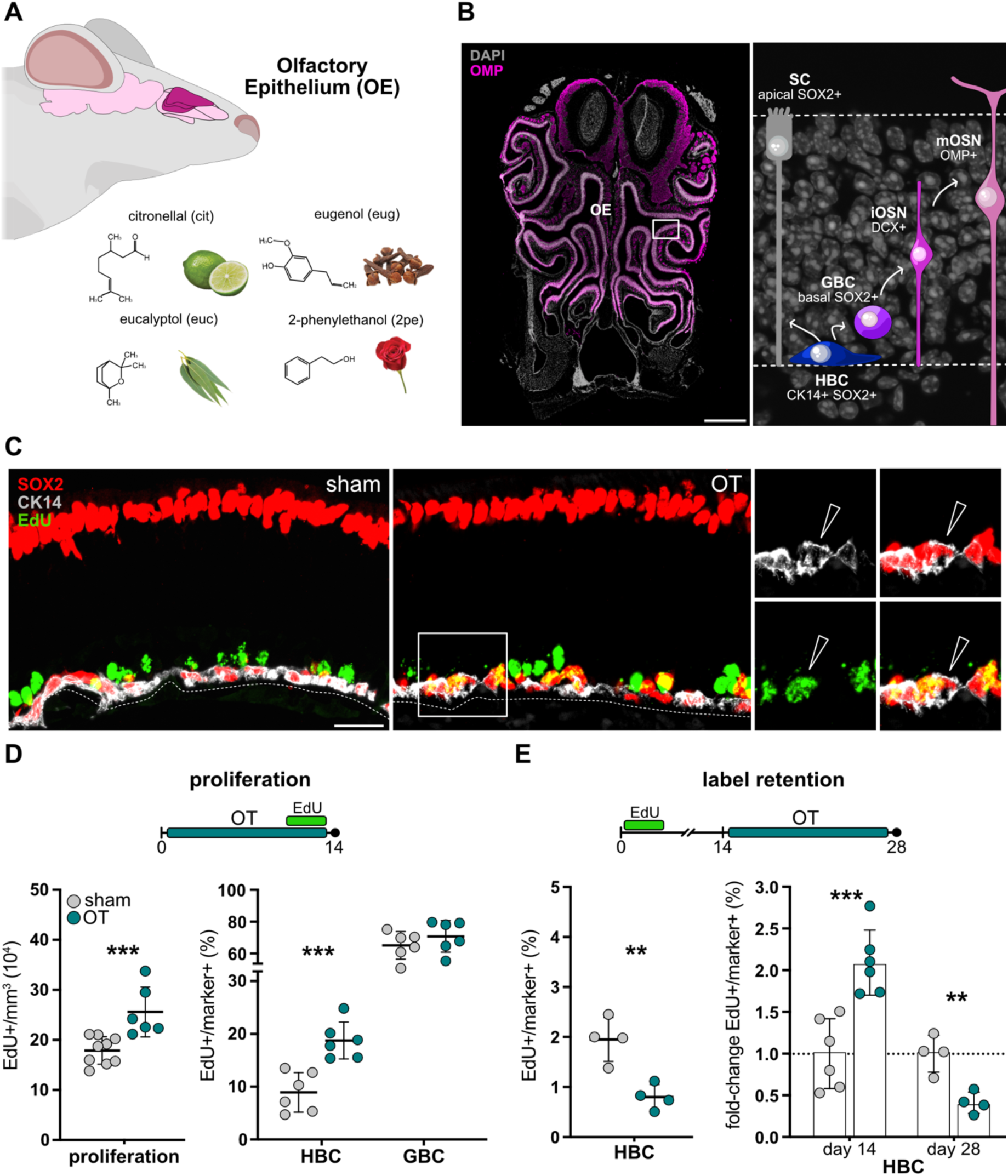
OT induces cell cycle re-entry of quiescent HBC. **A)** Cartoon of the mouse olfactory system highlighting the olfactory epithelium (OE, dark pink) and the odorants used during OT. **B)** Coronal section of the mouse OE (left) and schematic of cell lineages and markers used in this study (right) to identify HBC, GBC, immature and mature neurons (i- and m-OSN, respectively) and sustentacular cells (SC). **C)** Immunofluorescence pictures of the OE of sham and OT-treated mice used to identify EdU+ HBC (arrows in inset). **D** and **E)** Experimental protocols (top) and quantifications (mean ± SD; bottom) of proliferation and label-retention within the OE of sham or OT-treated mice (grey and blue dots, respectively). Cells counted per data point = *n* > 200. * = p ≤ 0.05, ** = p ≤ 0.01, *** = p ≤ 0.001. Scale bars = 200 (B) and 20 (C) μm.

To assess proliferation, mice were administered 5-ethynyl-2′-deoxyuridine (EdU) for 4 days prior to the end of OT (Fig. 1D; top). This revealed a nearly 50% increase in EdU+ cells in OT-treated mice compared with controls subjected to sham OT, in which water replaced the four odorants (sham vs. OT: 18.0 ± 2.7 and 25.7 ± 4.9·10^4^/mm^3^; *p* ≤ 0.001) (Fig. 1C and D; left). Next, we discriminated effects on proliferation within HBC (SOX2+/CK14+) and GBC (basal SOX2+) finding that OT more than doubled the proportion of EdU+ HBC (sham vs. OT: 9.0 ± 3.7 and 18.8 ± 3.5%; *p* ≤ 0.001) while no significant change was found among GBC (sham vs. OT: 65.2 ± 8.6 and 70.8 ± 9.9%; *p* = 0.32) (Fig. 1D; right).

The latter result was not surprising given that GBC under homeostatic conditions already have a high cycling activity^21^. Conversely, knowing that most HBC are quiescent^29^, we hypothesized that OT promoted their cell cycle re-entry. To test this, we designed a label-retention protocol in which mice were initially exposed to EdU for 4 days while starting OT 2 weeks later for additional 14 days (Fig. 1E; top). We argued that, under these conditions, cell cycle re-entry would be reflected by a decrease in label-retaining HBC. Consistently, at the end of OT, i.e. day 28, we found that EdU+ HBC were decreased by 50% in OT-treated mice relative to sham controls (sham vs. OT: 1.9 ± 0.4 and 0.8 ± 0.2; *p* ≤ 0.005) (Fig. 1E; left).

Systemic administration of EdU allowed us to assess proliferation also within the central neurogenic niche of the SVZ (Fig. S1A; top). Brains of mice used in the experiments above were processed for immunohistochemistry and stem/progenitor cells identified (SOX2+/S100β–). No statistical difference was found neither in overall proliferative activity (sham vs. OT: 6.5 ± 2.0 and 7.2 ± 2.5·10^2^/mm^3^; *p* = 0.68) nor proportion of EdU+ cells among neural progenitors (sham vs. OT: 49.0 ± 8.8 and 43.5 ± 12 %; *p* = 0.51) (Fig. S1A; bottom) indicating that OT induces proliferation specifically in the peripheral, but not central, olfactory system.

Finally, knowing that OT preserves its beneficial effects also in the elderly population^30^, we compared OE proliferation after olfactory stimulation in 6, 12 and 24 months old mice (Fig. S1B; top). Interestingly, while the proportion of proliferating HBC decreased during aging, OT remained similarly effective with 1.5 to 2-fold increases in the proportion of EdU+ HBC across ages (sham vs. OT fold-change: 2.3 ± 0.4, 1.9 ± 0.2 and 1.6 ± 0.2, for 6-, 12- and 24-months old mice, respectively) (Fig. S1B; bottom).

Collectively, our findings showed the cell-type specific effect of OT in promoting cell cycle re-entry of HBC. Intriguingly, effects on proliferation and label-retention essentially matched with a doubling in the former and halving in the latter (sham vs. OT fold-change: 2.1 ± 0.2 and 0.4 ± 0.1, respectively; all *p* ≤ 0.005) (Fig. 1E; right) suggesting that cell cycle re-entry is the most prominent, if not perhaps the only, effect of OT on quiescent HBC. These effects were specific to the OE of the peripheral nervous system, and nearly constant over the course of life, with no evident change at the level of the brain’s neurogenic niche of the SVZ.

### OT promotes the generation and survival of OSN

Next, we evaluated whether the increased proliferation of HBC triggered by OT correlated with increased neurogenesis. To this aim, we birthdated newborn neurons generated during 4 days of EdU administration at the end of OT and quantified them either immediately after the treatment, i.e. day 14, or two or four additional weeks later, i.e. day 28 or 42 (Fig. 2A; top). To account for their maturation, EdU+ neurons at day 14 were identified as immature OSN (iOSN) as assessed by immunoreactivity for DCX, which is expressed as early as 4 days after mitosis^21,31^ (Fig. 2A; left). Similarly, EdU+ mature OSN (mOSN) at day 28 and 42 were identified by OMP expression, which peaks 8 days after birth^32^ (Fig. 2A; right).

**Figure 2.**
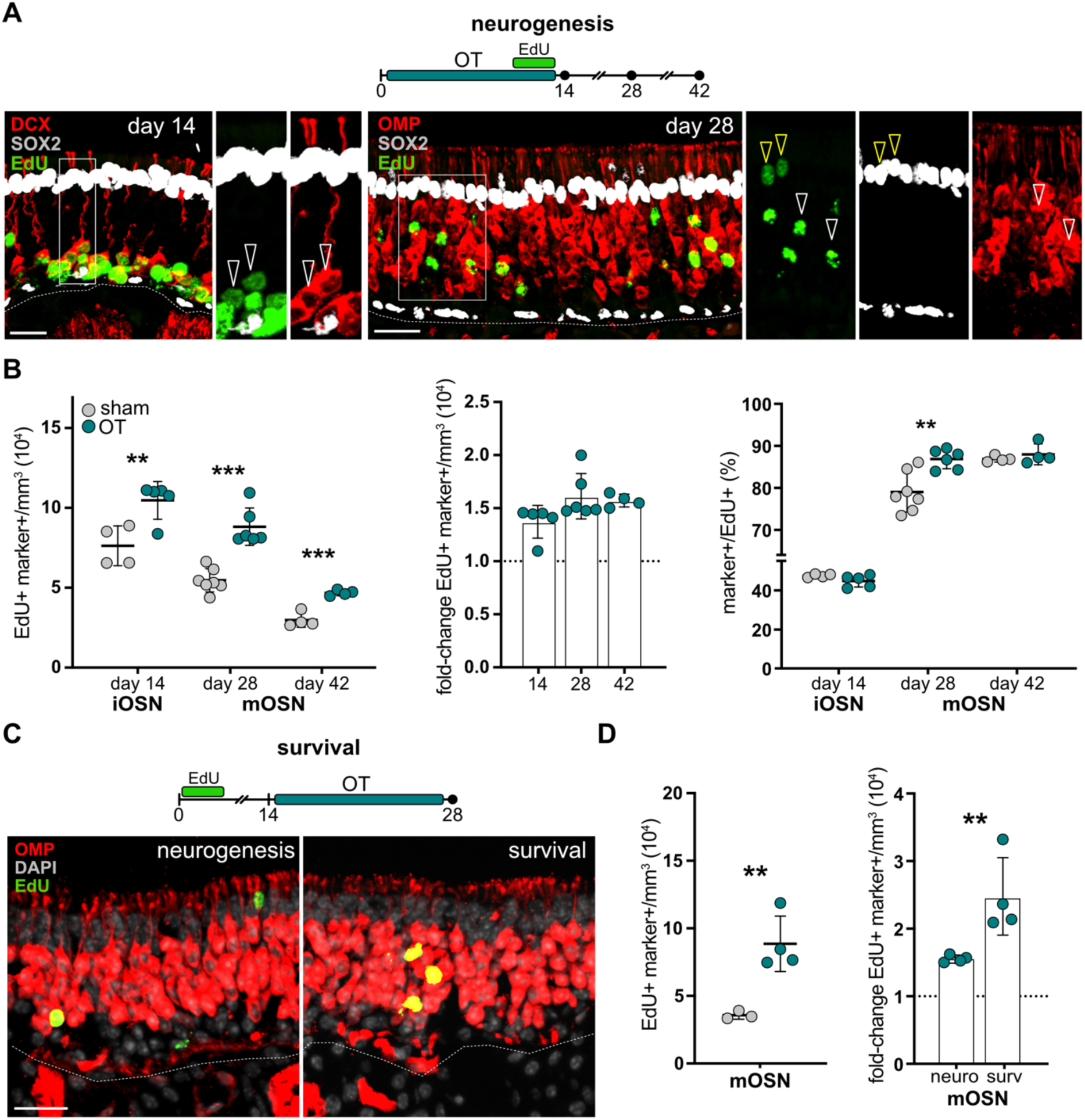
OT promotes generation and survival of OSN. **A)** OT experimental protocol to assess neurogenesis (top), immunofluorescence pictures of the OE (markers as indicated) combined with EdU labeling (arrows in insets = EdU+ cells) at day 14 or 28 (bottom) and **B)** quantifications of i/mOSN (mean ± SD). Total amounts, fold-changes and proportions of EdU+ OSN of control vs. OT-treated mice are depicted over time (left to right). **C)** OT experimental protocol to assess neuronal survival (top), fluorescence pictures of the OE (as in A) and **D)** quantifications (mean ± SD) of mOSN and fold-change effects between sham or OT-treated mice (grey and blue dots, respectively). Cells counted per data point = *n* > 250. * = p ≤ 0.05, ** = p ≤ 0.01, *** = p ≤ 0.001. Scale bars = 20 μm.

With regard to EdU+ iOSN, we found that their abundance at day 14 was increased by about 30% in mice subjected to OT relative to controls (sham vs. OT: 7.6 ± 1.2 and 10.4 ± 1.1·10^4^/mm^3^; *p* ≤ 0.01) (Fig. 2B; left). Two weeks later, at day 28, most EdU+ neurons effectively differentiated into mOSN, as confirmed by OMP immunoreactivity. Notably, the increased abundance of EdU+ iOSN observed at day 14 in mice subjected to OT persisted at later timepoints with equivalent increases in EdU+ mOSN both at day 28 (sham vs. OT: 5.4 ± 0.7 and 8.8 ± 1.1·10^4^/mm^3^; *p* ≤ 0.001) and day 42 (sham vs. OT: 3.0 ± 0.4 and 4.7 ± 0.2·10^4^/mm^3^; *p* ≤ 0.001) (Fig. 2B; left). In fact, while the abundance of birthdated EdU+ OSN decreased over the three time points, as expected based on their reported turnover^33^, the fold-change increase in OT-treated mice remained essentially the same (sham vs. OT, fold-change: 1.4 ± 0.1, 1.6 ± 0.2 and 1.6 ± 0.1 for day 14, 28 and 42, respectively) (Fig. 2B; center).

In addition to OSN, multipotent HBC generate all cell types of the mature OE. Hence, we next investigated whether OT triggered changes in the proportion of newly born OSN relative to non-neuronal cells (i.e. DCX–/OMP–). To this aim, we further classified EdU+ cells based on basal or apical tissue localization as, bona fide, label-retaining HBC or sustentacular cells, respectively. Interestingly, the proportion of OSN relative to non-neuronal EdU+ cells in OT-treated mice showed a transitory increase at day 28 (sham vs. OT: 79.0 ± 5.1 and 86.9 ± 2.3%; *p* ≤ 0.005), but not at day 14 or 42 (Fig. 2B; right and Fig.S2C). This transitory increase in OSN at day 28 was paralleled by a decrease in both basal and apical EdU+ cells (sham vs. OT: 11.2 ± 2.3 and 7.6 ± 1.5%; 9.7 ± 3.1 and 5.5 ± 0.9%, respectively, all *p* ≤ 0.01) (Fig. 2B; right and Fig.S2C). Yet, a physiological balance in the three EdU+ cell types was re-established at day 42 indicating that the increase in HBC proliferation triggered by OT (Fig. 1D) ultimately leads to an increase in all OE cell types in similar proportions. We therefore conclude that the transient over-representation of mOSN at day 28 likely reflects positive effects on neuronal maturation, as opposed to fate bias, following olfactory stimulation. These findings show not only that OT promotes neurogenesis but also that supernumerary OSN can mature and integrate into the OE according to their physiological turnover and fate.

However, the previous experiments could not address whether olfactory stimulation promotes survival of OSN generated prior to, rather than during, OT. To investigate this, we capitalized on our previous label-retention experiment in which EdU labelling was performed 2 weeks before OT, as opposed to at its end (Fig. 2C; top). At the end of this paradigm, i.e. day 28, we found that the abundance of EdU+ mOSN of OT-treated mice was more than doubled compared to controls (sham vs. OT: 3.6 ± 0.3 and 8.9 ± 2.1·10^4^/mm^3^; *p* ≤ 0.01) (Fig. 2D; left). This, in turn, indicated that olfactory stimulation promotes survival of mature neurons, in addition to and independently from, HBC proliferation, neurogenesis and OSN maturation.

More specifically, when comparing the relative contributions of OT to neurogenesis vs. neuronal survival (Fig. 2B vs. D) we found that the magnitude of the latter was nearly twice as much as the former (sham vs. OT fold-change: 1.6 ± 0.1 and 2.5 ± 0.6 for neurogenesis and survival respectively; *p* ≤ 0.01) (Fig. 2D; right).

As above, we also investigated whether effects of OT were specific for the peripheral, but not central, olfactory system and preserved during aging (Fig. S2A and B). With regard to the former, no difference was found in the abundance of the EdU+ DCX+ newborn neurons within the OB (sham vs. OT: 1.6 ± 0.2 and 2.0 ± 0.5·10^3^/mm^3^; *p* = 0.18) (Fig. S2A). With regard to the latter, and beside the expected age-dependent decline in EdU+ OSN, we found that OT increased neurogenesis by about 50% relative to age-matched controls across all ages analysed (sham vs. OT fold-change: 1.6±0.2, 1.4±0.2 and 1.5±0.4 for 6, 12 and 24 months old mice, respectively) (Fig. S2B).

Altogether, our data highlight the life-long effects of OT in promoting neurogenesis, neuronal maturation and survival within the OE, with no effect at the level the SVZ/OB. These effects were maintained throughout life such that, intriguingly, two years old mice undergoing OT rejuvenated to levels of neurogenesis comparable to that of 6 months controls (Fig. S2B).

### OSN activity recapitulates the effects of OT

While OT is beneficial in patients with olfactory dysfunctions, it is unclear why the plethora of odors that we perceive every day is insufficient to trigger similar effects. As one key difference between OT and natural stimuli, the former is performed with undiluted monomolecular odorants orders of magnitude more concentrated than natural odorants. We therefore investigated if the effects of OT were concentration-dependent by reproducing the experiments above (Fig. 1D, 2A and B) with odorants diluted 1:100.

We found that OT increased i) HBC proliferation, ii) neurogenesis and iii) neuronal survival also when using diluted odorants (sham vs. OT 1:100, fold-change: 1.6 ± 0.1, 1.2 ± 0.2 and 1.2 ± 0.1, respectively; all *p* ≤ 0.05), but at lower magnitudes compared to pure odorants (sham vs. OT pure fold-change: 2.3 ± 0.4, 1.6 ± 0.2 and 2.5 ± 0.6 for proliferation, neurogenesis and survival, respectively; all *p* ≤ 0.05) (Fig. 3A and B). This explains not only why odorants at natural concentrations are less effective in patients but also suggests that the activity and/or proportion of OSN stimulated by odorants is the main trigger of HBC proliferation and neurogenesis.

**Figure 3.**
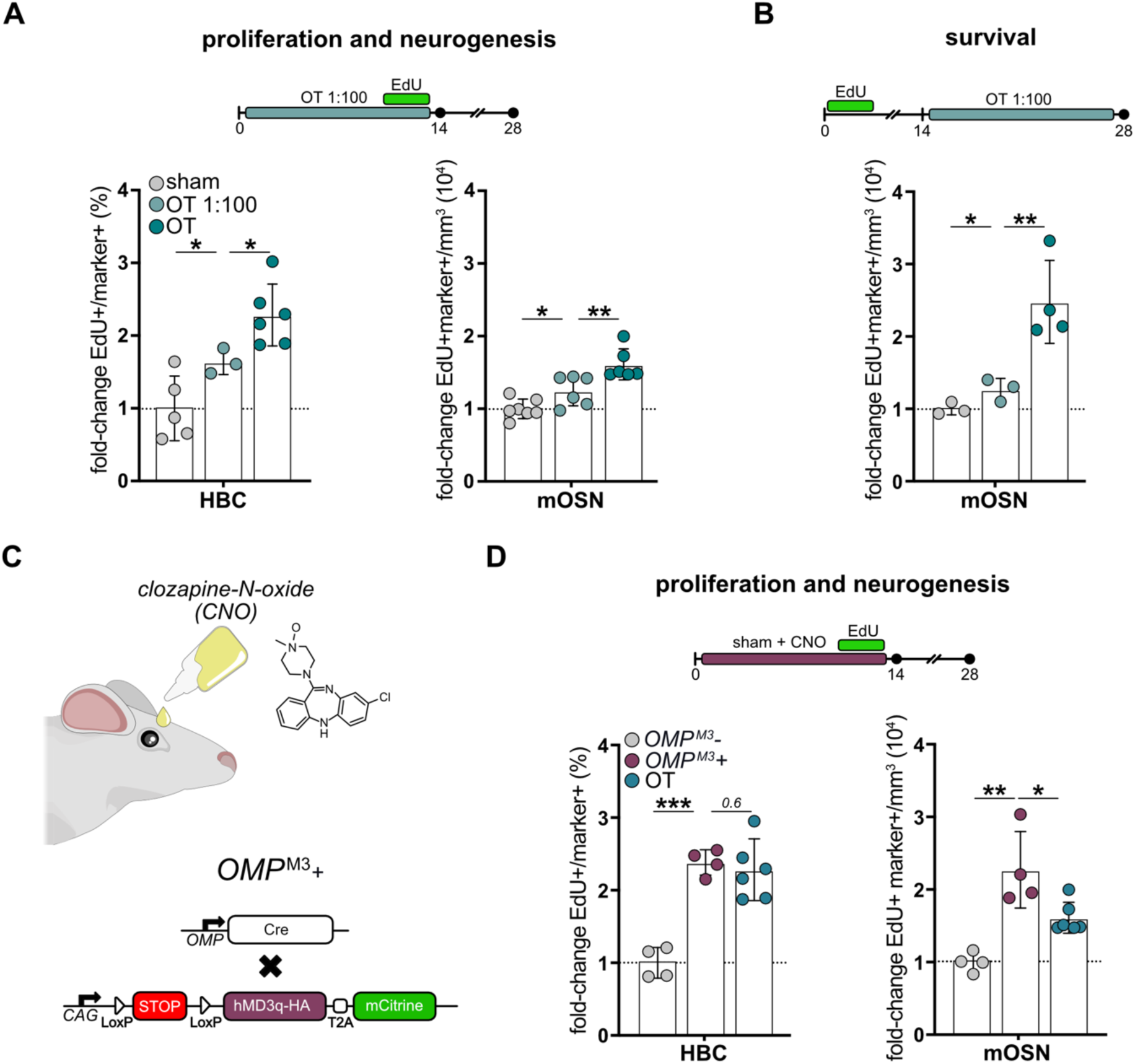
OSN activity alone recapitulates the effects of OT. **A, B** and **D)** OT experimental protocols (top) and fold-change quantifications (bottom; mean ± SD) of proliferation, neurogenesis and neuronal survival of wilt-type mice subjected to OT with 1:100 diluted odorants (light blue dots in a and b) or *OMP*^M3^+ and OMP ^M^^3^*–* mice (purple dots and grey dots, respectively, in D. **C**) Genetic design of *OMP*^M^^3^ mice with CNO administration by eye drops. Data showing the fold-change effects of OT with pure odorants (A, B and D; dark blue dots) and sham (in A and B; grey dots) were taken from previous experiments (Fig. 1 and 2). Cells counted per data point = *n* > 200. * = p ≤ 0.05, ** = p ≤ 0.01, *** = p ≤ 0.001.

The effect of OT on proliferation of HBC was in fact intriguing. While OSN maturation and survival are expected to be promoted by neuronal activity, HBC are not directly exposed to the odorants themselves and are not expected to express olfactory receptors. Hence, we next tested whether OSN activity alone was sufficient to trigger HBC activation in the absence of olfactory stimulation.

To this aim, we generated an OSN-specific excitatory DREADD mouse line, allowing us to activate OSN by CNO (*OMP^M3^+* line, see methods; Fig. 3C). Efficacy and specificity of DREADD expression in *OMP*^M3^+ mice and *OMP*^M^^3^*–* littermates negative controls were validated by immunohistochemistry and co-localization of the DREADD-reporter (mCitrine) together with OMP. While no reporter expression could be observed in control *OMP*^M^^3^*–* mice, an almost complete overlap between mCitrine and OMP was found in *OMP*^M3^*+* littermates (Fig. S3A). In essence, the *OMP*^M3^+ line allowed us to excite virtually all OSN selectively with CNO in the absence of olfactory stimulation.

*OMP^M3^+* mice and *OMP^M^*^3^*–* controls were subjected to sham OT, with both receiving CNO twice per day for 14 days and EdU being administered for 4 days as above (Fig. 1D and 2A). Mice were sacrificed on day 14 or 28 to assess HBC proliferation and neurogenesis, respectively. At the first timepoint, i.e. day 14, the proportion of EdU+ HBC in *OMP^M3^+* mice increased relative to littermate controls (*OMP^M^*^3^*–* vs *OMP^M3^+*fold-change: 2.4 ± 0.2; *p* ≤ 0.001) to a nearly identical level than that of wildtype mice subjected to OT (Fig. 3D; left). At the second timepoint, i.e. day 28, also the density of EdU+ mOSN in *OMP*^M3^*+* mice increased (*OMP^M^*^3^*–* vs. *OMP^M3^+* fold-change: 2.3 ± 0.5; *p* ≤ 0.005). Notably, this latter effect was even greater that that observed in OT-treated animals (sham vs OT fold-change: 1.6 ± 0.2; *p* ≤ 0.05) (Fig. 3D).

Collectively, the concentration-dependent effects of OT explain why natural odorants are less beneficial to patients, while their replication upon OSN activation alone provides insights for alternative interventions for OT-unresponsive patients.

### OT improves odor discrimination by mitral cells

While OT promotes the generation and survival of OSN, it is unclear whether and how increased OE neurogenesis improves odor coding in downstream olfactory circuits. To assess odor representations in the OB, we next performed time-lapse, two-photon calcium imaging in head-fixed, awake mice expressing GCaMP6s selectively in MC (Fig. 4A and B). First, we imaged responses of MC to the odorants used during OT plus an additional, fifth odorant, isoamylacetate (iso), as a generalized stimulus. Next, mice were subjected to OT and two weeks later re-imaged after stimulation with the same five odorants. We identified and analyzed a total of 95 MC recorded before (T1) and after (T2) OT (n = 4 mice).

**Figure 4.**
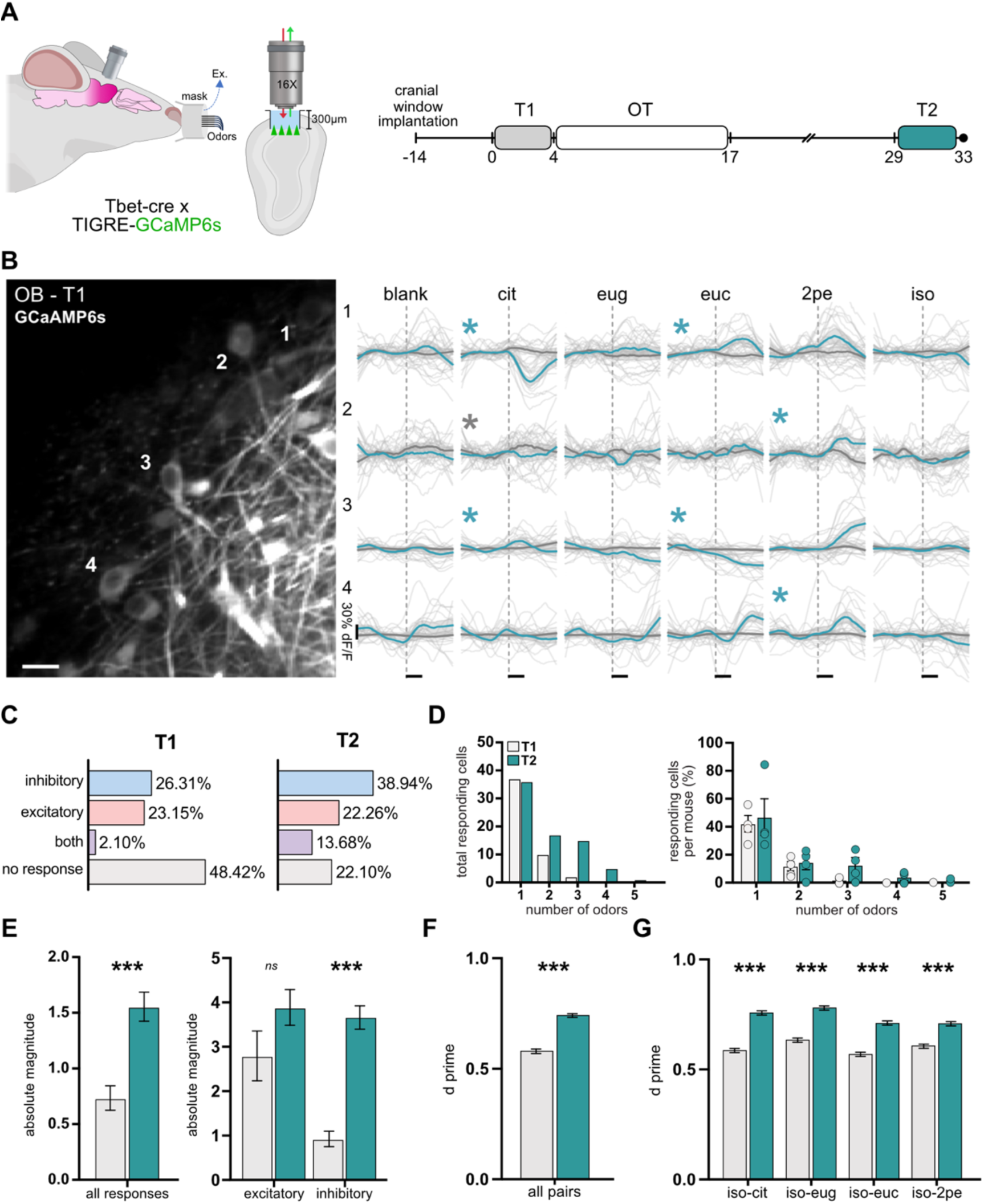
OT enhances MC discriminability in the OB. **A**) Cartoon illustrating two-photon calcium imaging of GCaMP6s-expressing MC (left) and experimental protocol to assess MC responses before (T1) and after (T2) OT (right). **B**) Example of a 2-photon micrograph of MC (left) and their representative calcium traces to the five odorants at T1 and T2 (left; gray and blue, respectively, thin lines = single trials, thick lines = average responses, horizontal scalebar represent stimulus). **C**) Proportion of MC showing inhibitory, excitatory, both, or non-responsive. **D**) Distribution of number of odors evoking MC responses for total number of cells (left) and averaged per mouse (right). **E**) Magnitude of inhibitory and excitatory responses pooled together (left) or separated (right). **F**) Pairwise *d′* odor discriminability values at T1 and T2 (gray or blue, respectively) per odor pair. **G**) Population-level *d′* values across isoamylacetate pairs. * = p ≤ 0.05, ** = p ≤ 0.01, *** = p ≤ 0.001. Scale bar = 50 µm.

Consistent with previous reports^34–36^, MC responses were heterogeneous, encompassing both excitatory and inhibitory, odor-evoked activity (Fig. 4C and D). In our analysis, the mean population responses from all MC–odor pairs did not significantly change after OT (Fig. S4A and B, cell mean). This was not surprising considering the diversity of odor qualities tested. In contrast, when analyzing single-neuron responses to one or more odor stimuli, a number of differences were observed between T1 and T2. Not only the number of MC unresponsive to all 5 odorants decreased at T2 relative to T1 (from 48 to 22%) but also among T1 responsive cells, their responses at T2 switched from inhibitory to excitatory, vice versa, or both (Fig. 4C and S4C). Overall, OT was associated with a greater proportion of MC exhibiting inhibitory responses (T1 vs. T2: from 26 to 39%) as well as an increase in cells showing both inhibitory and excitatory activity, depending on the odorant used (T1 vs. T2: from 3 to 14%) (Fig. 4C).

Moreover, we quantified tuning breadth by counting, for each MC, how many of the five odorants (0–5) evoked a significant response before and after OT (Fig. 4D). This showed that the proportion of completely unresponsive MCs was reduced by half upon OT (T1 vs T2: from 48 to 22%) while, at the same time, the fraction of cells responding to multiple odorants increased by three fold. For example, from T1 to T2, the number of MC responsive to 2, 3, or more odorants increased from 10, 2 and 0 to 17, 15 and 5, respectively (Fig. 4D, left). Paired analysis across the same 95 tracked MCs showed that OT increased the number of odorants driving a response (Fig. 4D). When tuning breadth was averaged per mouse, three out of four animals showed an increase after OT, although this trend did not reach significance at the mouse level (Fig. 4D, right).

In addition to changes in the proportion of MC responding to odorants, and more prominently, the magnitude of their responses as assessed by intensity of GCaMP6s fluorescence was markedly enhanced, both at the population level (T1 vs. T2: 0.7 ± 0.1 and 1.5 ± 0.1 absolute magnitude; *p* ≤ 0.001) (Fig. 4E) and across most individual odors (Fig. S4D and E). Notably, this increased inhibition was associated with higher odor discriminability, as quantified by pairwise *d′* analysis, for all odor pairs (T1 vs. T2: 0.5 ± 0.0 and 1.7 ± 0.0; *p* ≤ 0.001) (Fig. 4F). Interestingly, an increased discriminability was also observed when considering pairs including the fifth generalized stimulus, isoamylacetate (Fig. 4G), suggesting that the effects of OT are not limited to the odorants used during treatment.

These results indicate that OT improves MC sensitivity and discriminability, primarily through increased inhibition.

## DISCUSSION

To this day, OT remains the most effective clinical treatment for olfactory dysfunctions affecting one in four people worldwide^11^. Since its first description^10^, OT has become a common medical practice in many countries. Yet, the mechanisms mediating its efficacy remain elusive. Understanding the biology underlying OT is important to help define more effective therapies for the nearly half of the patients unresponsive to the treatment. By modeling OT in mice, here we show that olfactory stimulation triggers stem cell proliferation, neurogenesis, and OSN survival. Most importantly, OT refines odor processing in the OB increasing MC sensitivity and discriminability. Several aspects of our study are worth discussing and opening up avenues for future research.

First, while many studies have long reported the effects of physiological stimuli, such as physical or cognitive tasks, on adult neurogenesis in the brain^37,38^, to the best of our knowledge our is the first report highlighting a similar effect in the peripheral nervous system. In this context, we show that neuronal activity alone is sufficient to induce cell cycle re-entry of quiescent HBC, ultimately resulting in increased neurogenesis. Consistent with our finding, activity of granule cells in the adult hippocampus was found to regulate stem cell quiescence by signaling pathways requiring cell-cell contact^39^. Whether or not the same mechanism holds true for HBC remains to be investigated but, irrespective of this, our finding suggests that direct electric stimulation of the human olfactory mucosa may aid regeneration in a clinical setting.

Second, our study reveals the dose-dependent effect of olfactory stimulation on neurogenesis and neuronal survival. Such effects were significantly reduced when using diluted odorants and, in a series of experiments using single odorants, barely noticeable (not shown). This not only explains why patients who are normally exposed to everyday smells often show lower frequencies of olfactory recovery but also suggests that the nature and intensity of the stimuli are critical to achieve functional regeneration. Notably, while the four odorants originally used for OT were chosen to encompass a broad spectrum of odor qualities, odor perception is characterized by a high degree of individuality due to extensive genetic variation of olfactory receptors and life style^40,41^. In other words, providing insight into future personalized medicine, our study suggests that OT regimens designed on patient’s olfactory perception, and receptors expression, will be more effective.

Finally, with no neurogenic effect within the SVZ-OB, our study suggests that OT mainly acts at the level of OSN–MC connectivity. This raises an important question with regard to how neurogenesis in the OE influences odor-evoked activity in the OB. Although we did not directly measure connectivity between the OE and OB, our functional imaging data support a model whereby an expanded pool of OSN form additional synapses onto MC, thereby boosting their afferent drive to MC. This interpretation is consistent with the broadening of MC tuning to more odorants, that we observed at the single-cell level after OT, including the recruitment of additional inhibitory responses. Although the corresponding mouse-averaged measure only showed a trend with our sample size, the direction of the change was consistent in three out of four animals.

Concomitantly, the same increase in OE input is expected to recruit activity-dependent inhibitory pathways in the OB^16,42,43^,overall strengthening inhibition and supporting odor discriminability. The changes in response sign and tuning of MC following OT may seem to be conflicting with better discriminability. However, the heterogenous nature of odor coding in the OB could explain both observations. For example, changing one odor response from weak excitation into net inhibition, while another remains excitatory, can result in better response differences despite wider tuning, ultimately explaining both broader responsiveness and enhanced discriminability upon OT. Notably, these effects held true for trained as well as a novel odor, which is fully consistent with a broad clinical literature showing that OT improves odor sensitivity, discrimination, and identification across all odorants, not only those used during OT^11^. While an increased synaptic drive from OSN-to-MC provides one conservative account of our results, other processes may well contribute. For example, shifts in the balance among local inhibitory circuits and OSN excitation^16,44^ , and long-range cortical feedback ^45,46^ that may all modulate the observed plasticity following OT and merit future investigation.

## LIMITATIONS OF THE STUDY

Our study was limited to mice and could not directly assess humans. Yet, the biology of the sense of smell is highly conserved across mammals^1^ making it likely that our fundamental observations also apply to humans. It is important to note, however, key differences between our study and the clinical treatment. For a start, contrary to patients, we used mice with normal olfactory function whose OE was not damaged by infections or lesions. In addition, OT in mice was limited to 2 weeks, significantly shorter than the 3 months usually adopted as clinical treatment. Finally, proportion of cell types, cell-cycle kinetics, and turnover may also contribute important differences across species. Yet, a regenerative context and longer treatment in patients seem to concur into making effects in humans greater than we describe in mice. Particularly for prolonged treatments, effects on neurogenesis and neuronal survival may act synergistically thereby an increased number of neurons would feedback with their activity to further promote HBC expansion.

## RESOURCE AVAILABILITY

All relevant data are available from the authors upon request.

## ACKNOWLEDGEMENTS

This work was supported by a German-Israeli DFG research grant (CA 893/22-1), the CRTD of the TU-Dresden. The work was supported by the Gatsby Charitable Foundation. E.T. was supported by a postdoctoral fellowship from the Edmond and Lily Safra Center for Brain Sciences; A.M. is the Eric Roland Chair in Brain Sciences. We thank Meirav Givon and Joseph Zak for Technical help and discussion, respectively. We thank Idan Segev for advice and support on this project.

## AUTHOR CONTRIBUTIONS

V.A., T.H., A.M. and F.C. conceived and designed the study. V.A., E.T. and G.K. performed the experiments. V.A., E.T. and S.S. analyzed the data. V.A., E.T., A.M and F.C. designed the figures and wrote the manuscript. F.C., S.I. and A.M. obtained the funding. All the authors discussed the data and revised the final version of the manuscript.

## DECLARATION OF GENERATIVE AI and AI-ASSISTED TECHNOLOGIES

During the preparation of this work, the authors used ChatGPT and Grammarly only for editing purposes. After using this tool or service, the authors reviewed and edited the content as needed and take full responsibility for the content of the publication.

## SUPPLEMENTAL INFORMATION

***Document S1***. Figure S1-S4, Table S1-S2

## STAR METHODS

### Animals

Experiments were approved by local authorities (TVV-12/2021, 19/2024 and NS-22-16909-4) with mice kept in standard cages with a 12 hour light cycle and water and food ad libitum. Wildtype females (C57BL/6JRj) were used to assess the cellular effects of OT upon exposure to pure (or diluted 1:100) odorants in the following order: citronellal, eugenol, eucalyptol and 2-phenylethanol (Sigma; Table S1). Specifically, four pieces of paper towel were soaked with 500 μl of each odorant and placed in petri dishes covered with perforated parafilm to allow odor diffusion without contact. OT involved two daily exposures (morning and evening), 3 min each odorant, for 14 days. Water instead of odorants was used for sham OT. DREADD and GCaMP6s mouse lines were used for chemogenetic activation of OSN or MC imaging, respectively. The former was obtained by crossing *OMP::*Cre (Jax stock #006668) (Li et al., 2004) with *CAG::*STOP-flox-hM3Dq-mCitrine (Jax stock #026220) (Zhu et al., 2016) mice and OSN activity triggered by 6 μl of clozapine-N-oxide (Enzo Life Sciences GmbH) dissolved in PBS (1mg/kg) by eye-drops ^47^, 1 hour before sham OT. Littermates lacking Cre were used as negative controls. The latter mouse line was obtained by crossing Tbet-Cre mice (Jax stock #024507) ^48^with Ai162 mice (TIGER-GCaMP6s) (Jax stock #033361) ^49^ (both C57BL/6 genetic background) and used for calcium imaging (see below). EdU (Sigma-Aldrich) was dissolved in PBS and 100 μl administered intraperitoneally (5 mg/kg), twice daily, for 4 days. For histology, mice were anesthetized and transcardially perfused with saline, followed by fixation with 4% paraformaldehyde (PFA) in phosphate buffer.

### Two-photon Calcium imaging

Mice were anesthetized with ketamine and domitor (10 mg/kg, intraperitoneal) and carprofen (4 mg/kg, subcutaneous). Anesthesia was monitored via the pinch-withdrawal reflex and rectal temperature maintained at 36.5 ± 0.5°C. Upon incision of the head skin, a metal bar was affixed to the skull with dental cement and a 2.5-mm cranial window implanted over one OB secured with a triple-layered window (2 x 3 mm) as previously described ^50^. Imaging sessions began 7-10 days following cranial window implantation and mice habituated to the head-fixed setup for 2–4 days. OT odors plus isoamylacetate were initially diluted in mineral oil (100 ppm), mixed with air, and delivered (900 ml/min flow) using a six-odor air-dilution olfactometer (RP Metrix Scalable Olfactometer Module LASOM 2). Before reaching the final valve (via a four-way Teflon valve, NResearch), odors were further diluted tenfold (10 ppm). During stimulus, the final valve switched to direct the odorized airflow to the odor port and reaching the mouse nostrils via a custom-made glass mask at a flow rate of 1 l/min (2 s duration, 10 s interstimulus interval). Continuous air suction removed residual odors. During interstimulus intervals, a stream of filtered air (1 l/min) flowed to the odor port. The olfactometer was calibrated using a miniPID (Aurora Scientific). Imaging of the OB was performed using an Ultima two-photon microscope (Prairie Technologies) equipped with a 16X water-immersion objective (0.8 NA; CF175, Nikon). Two-photon excitation was delivered at 920 nm using an Axon laser (Coherent, United Kingdom), tuned to 920 nm for excitation of GCaMP6s. Emitted fluorescence was collected through a bandpass filter (525/50 nm) and detected by GaAsP photomultiplier tube. Image acquisition was carried out at a frame rate of 7 Hz with a resolution of 400 × 200 pixels per frame, using PrairieView software 5.5. Each imaging session consisted of 12 odor presentation trials per odor (7 s baseline, 2 s odor presentation and 5 s post-stimulus). Each mouse was imaged across 3–4 sessions lasting 45–60 min each. Between 3-4 fields of view were acquired per mouse, with each field being imaged across all odor trials. The same MC were imaged before and after OT and tracked across sessions, enabling within-cell paired comparisons of odor-evoked responses before and after training.

### Immunohistochemistry

Snouts, containing the OE, were decalcified with EDTA (0.5M, pH 7.0 in PBS) for 4 days and cryoprotected in sucrose-based solution (10 and 20% 1 h at room temperature and 30% overnight at 4°C). Snouts were next embedded in Tissue Tek O.C.T compound (Science services), snap-frozen in dry ice, cut into 12 μm thick cryosections along the antero-posterior axis and collected on Superfrost Plus adhesive microscope slides (Epredia – SuperfrostTM) and stored at –20°C. Brains, including were post-fixed overnight in 4% PFA and sectioned into 40 μm thick vibratome sections, stereologically collected across the SVZ or OB, and stored at −20°C in cryoprotectant solution (25% ethylene glycol and 25% glycerol in PBS). After blocking and permeabilization with 10% donkey serum in 0.3% Triton X-100 in PBS for 1.5 h at room temperature, primary and secondary antibodies (Supplemental Table 2) were incubated in 3% donkey serum in 0.3% Triton X-100 in PBS overnight at 4°C. EdU was detected following the manufacturer’s guidelines (Click-iT EdU, ThermoFisher Scientific) and DAPI used to counterstain nuclei.

### Cell quantifications and statistics

Fluorescence pictures were acquired using an automated Zeiss ApoTome (Zeiss). For the OE, a minimum of 3 sections were imaged per animal, corresponding to the anterior, medial, and posterior axis of the OE. Within each section, the dorsal turbinate was identified and 5 optical sections (10 mm thick, 2 mm interval) imaged and quantified. For SVZ and OB quantifications, 1 and 3 sections were imaged respectively, and 5 optical sections (20 mm thick, 4 mm interval) were acquired. Stacks of different focal planes or tile mosaics were acquired using the microscope optical sectioning system when required. The stitching of tile mosaics and the generation of maximal intensity projections of the optical section were performed using the ZEN software (Zeiss). Images were further processed with Affinity Photo (Serif), and cells of interest quantified as required. In density quantification, areas were measured with Fiji/ImageJ and volume calculated according to optical thickness. Quantified cells numbers (*n*) and number of animals (*N*) are indicated in respective figure legends. Data are reported as mean ± SD and statistical significance was calculated by two-tailed unpaired Student’s t test assuming equal variance.

### Calcium imaging analysis

Data were analyzed using Python 3.11 and MATLAB R2024b (MathWorks). Initial preprocessing and regions of interest (ROI) extraction were performed using Suite2p (https://suite2p.readthedocs.io/en/latest/), which enabled the automatic detection and segmentation of MC bodies. For each identified ROI, we extracted the raw fluorescence signal (F) and an associated local neuropil signal (Fneu). To correct for background fluorescence, the corrected signal was computed as: Fcorr = F – 0.7*Fneu. The relative fluorescence change (dF/F = (Fcorr – F0) / F0) was calculated, with baseline fluorescence (F0) defined as the median fluorescence over the 5 seconds preceding odor onset ((-5) – 0 s), for each cell in each trial. Traces were smoothed with a Savitzky-Golay filter (7 frames window). Odor-evoked responses were quantified by comparing mean signal in the 0-4 seconds post-stimulus interval, to the mean signal (-4) - 0 seconds pre-stimulus baseline, using a two-sided Wilcoxon signed-rank test. Only responses with a p-value ≤ 0.05 (using nan_policy=’omit’ to handle any missing data) were considered statistically significant. Responses were classified as inhibited or excited based on the sign of the integral of dF/F traces. Response magnitude was computed by subtracting the mean signal over the seven frames immediately preceding the stimulus onset and integrating the baseline-corrected trace via trapezoidal numerical integration over the 0–4 seconds post-stimulus interval. Temporal magnitude analysis was assessed by computing the area under the curve in predefined time windows. The proportion of responding cells per odor (1-5) was determined separately for excited and inhibited responses and statistical significance between T1 and T2 was calculated by Two-sample Kolmogorov-Smirnov test. To quantify tuning breadth (Fig. 4D), we counted for each mitral cell how many of the five odorants (0–5) evoked a significant response at T1 and at T2, separately for excitatory responses, inhibitory responses, and their union. Changes in tuning breadth were first assessed at the cell level using paired Wilcoxon signed-rank tests across the 95 tracked mitral cells (*n* = 95 cells from 4 mice). Because the mouse is the primary experimental unit, we also averaged tuning breadth per mouse and compared T1 and T2 using a Wilcoxon signed-rank test (*n* = 4 mice). Only cells with clear morphological profiles were included in the analysis; significant stimulus-evoked fluorescence responses in these cells were categorized as excitatory or inhibitory.

Discriminability (*d’*) was used to quantify the separation between population responses to odor pairs, computed as:

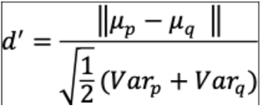

where p, q are responses to specific odors (with size of 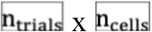). 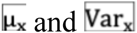 denote the mean and variance across trials for odor x (each a length 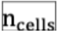 vector). The numerator is the Euclidean (L₂) distance between the two mean response vectors, and the denominator scales this distance by the average trial-to-trial variability of the two odors. Data were statistically analyzed using MATLAB, and Python with two-sided tests and are reported as mean ± SEM, unless stated otherwise.

