## Supplementary material for "Olfactory Training Enhances Adult Neurogenesis of the Olfactory Epithelium Improving Odor Discriminability of the Olfactory Bulb"

**SUPPLEMENTAL INFORMATION**

***Document S1***. Figure S1-S4, Table S1-S2

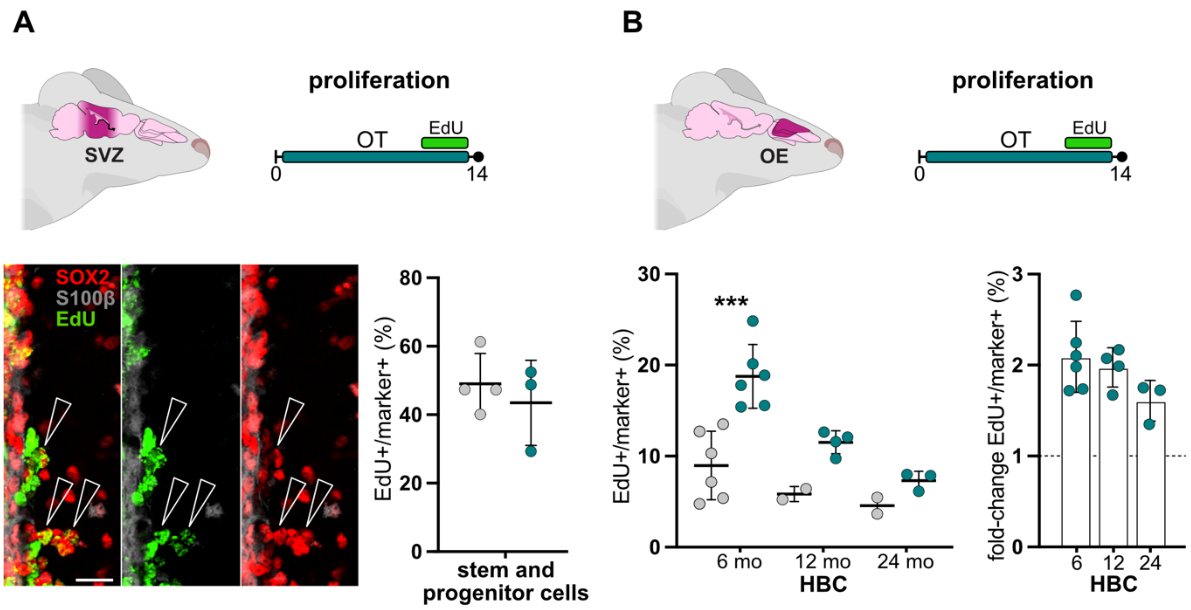


**Supplemental figure 1 – OT effects on proliferation in the SVZ or OE during aging A** and **B)** Schematic representation of neurogenic niches and OT experimental protocols (top; left and right, respectively), fluorescence pictures of the SVZ immunostained for markers of interest and EdU labelling (arrows = EdU+ progenitors) and quantifications (bottom; mean ± SD) of proliferation among neural progenitor cells (A; right) and among HBC during aging with respective fold-changes (B, left and right, respectively) in sham or OT-treated mice (grey and blue dots, respectively). Cells counted per data point = $n > 200$. * = $p \leq 0.05$, ** = $p \leq 0.01$, *** = $p \leq 0.001$. Scale bar = 20 µm.

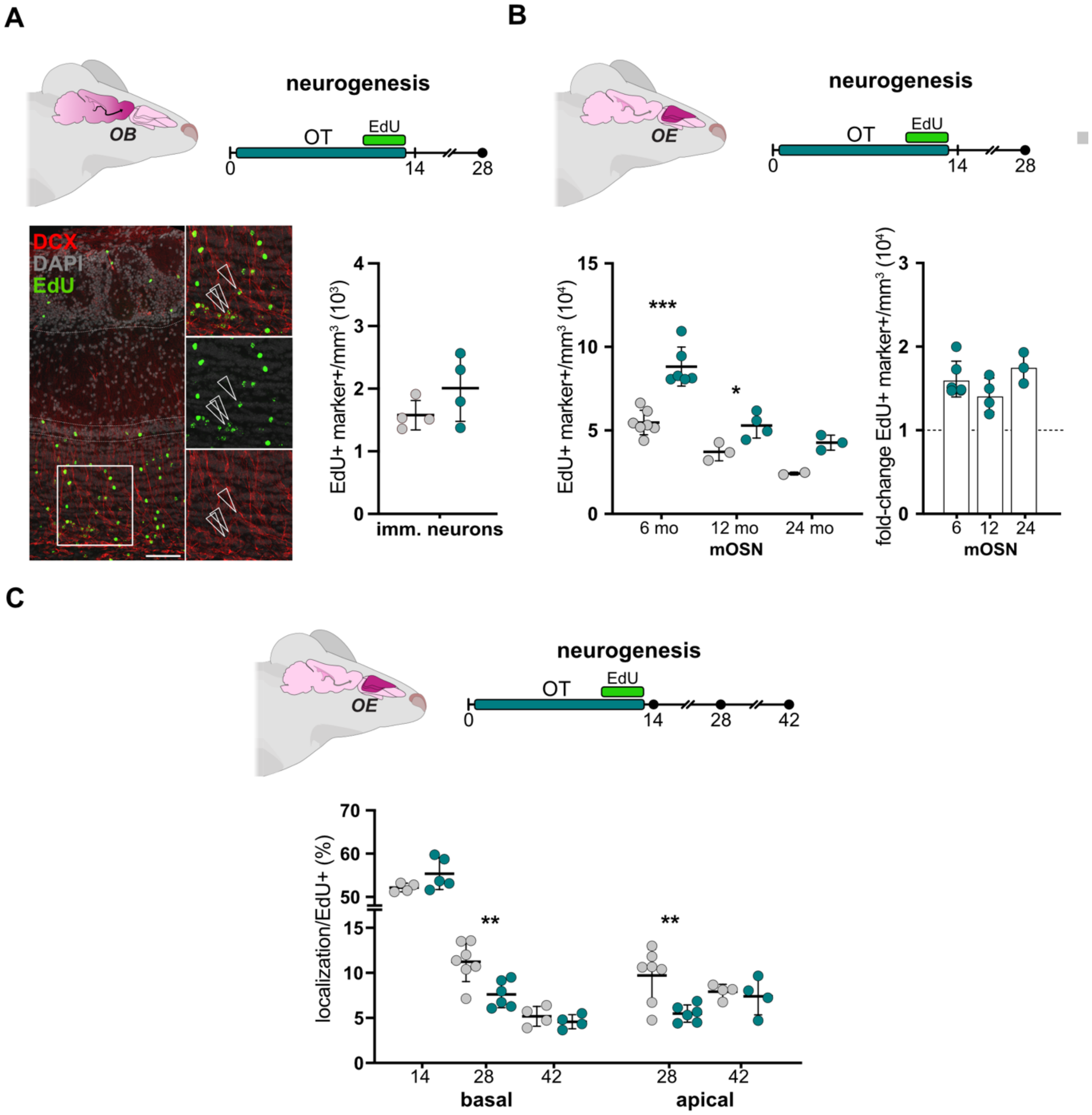


**Supplemental figure 2 – OT effects on neurogenesis in the OB and during aging A-C)** Cartoons depicting neurogenic niches and OT experimental protocols (top) used to quantify (bottom; mean ± SD) OB adult born neurons upon immunohistochemistry for markers of interest and EdU labelling across lifespan (B, left; arrows in inset) or EdU+ cells across boundaries of the OE (C; apical or basal) of sham or OT-treated mice (grey and blue dots, respectively). Cells counted per data point = n > 200. * = $p \leq 0.05$, ** = $p \leq 0.01$, *** = $p \leq 0.001$. Scale bar = 200 µm.

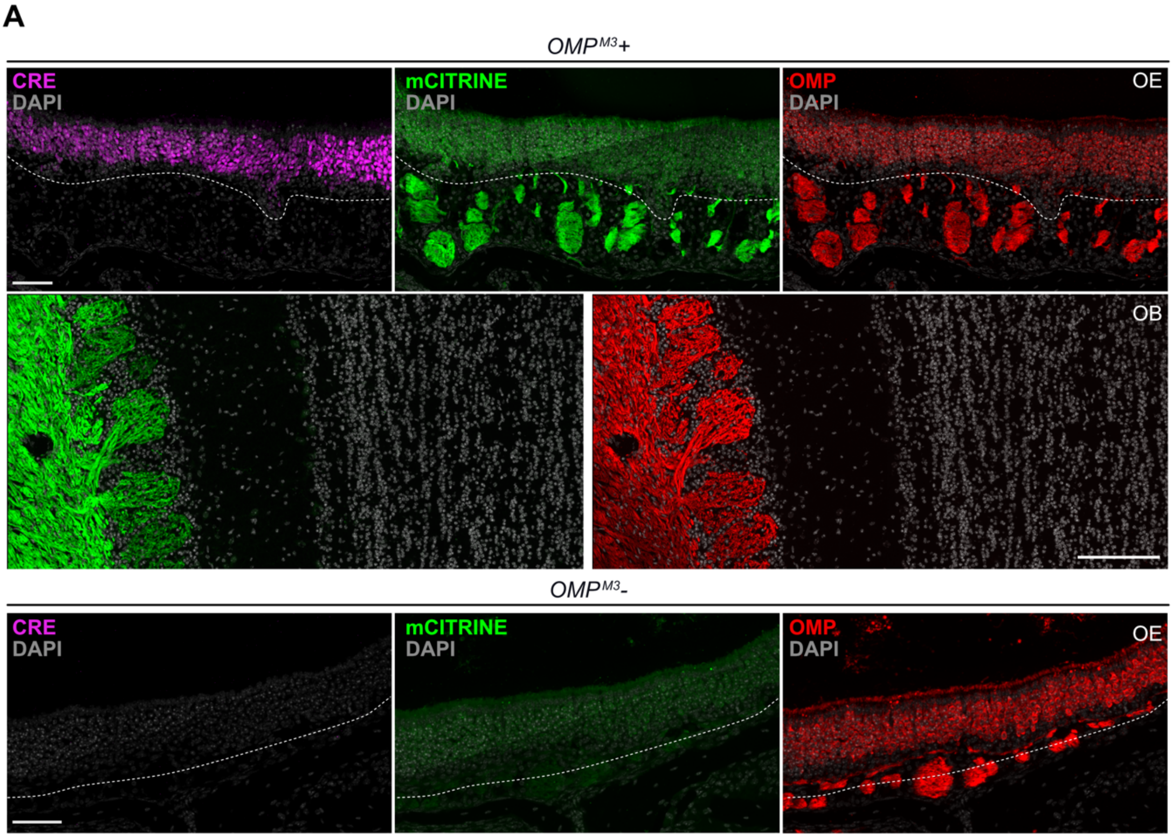


**Supplemental figure 3 – Effects of OSN excitation on proliferation and neurogenesis. A)** Fluorescence pictures of the OE and OB from *$OMP^{M3}$*+ (top) and of OE from *$OMP^{M3}$*– (bottom) immunostained for markers (as indicated) and DAPI counterstaining. Scale bar = 50 µm.

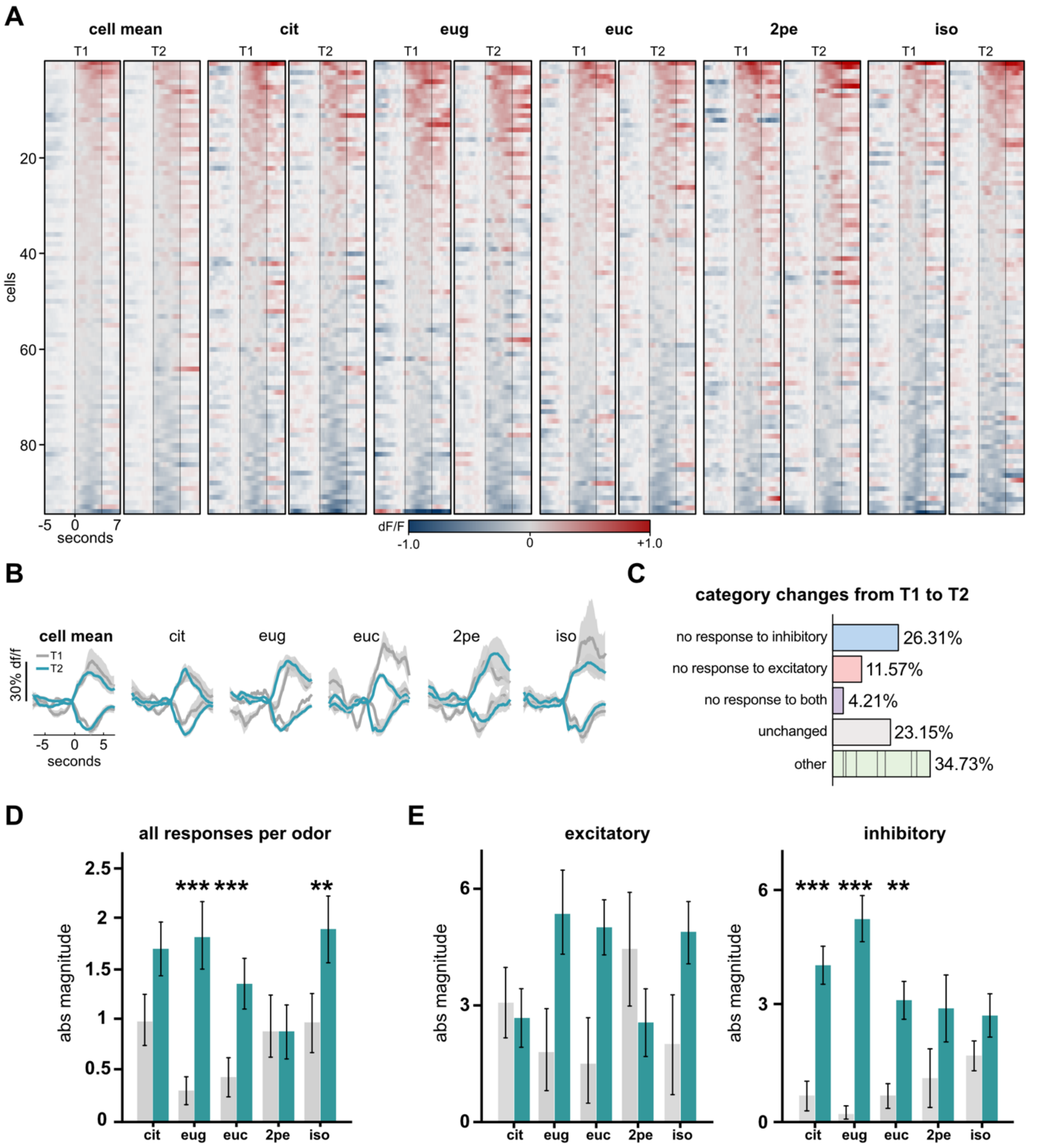


**Supplemental Figure 4 – Dataset of MC calcium responses A)** Heatmaps of normalized calcium responses for all MC–odor pairs before and after OT. Cells are sorted from most excited to most suppressed, based on the "cell's mean" response (left columns). **B)** Average population responses per odor before vs. after OT. **C)** Categorization of MC responses before and after OT showing most relevant switches in neuron response type. Additional switch combinations, or lack thereof, were pulled together (others). **D** and **E**) Overall absolute response magnitudes before vs. after OT, pooled across cells and presented per odor. * = p ≤ 0.05, ** = p ≤ 0.01, *** = p ≤ 0.001.

**Supplemental Table 1 – Odorants used in this study**

| Odorant | Molecule | Catalog No. | Supplier |
|---|---|---|---|
| citronellal (cit) | $C_{10}H_{1}3O$ | 27470-100ML-F | Sigma |
| eugenol (eug) | $C_{10}H_{12}O_{2}$ | W246719-1KG-K | Sigma |
| eucalyptol (euc) | $C_{10}H_{18}O$ | C80601-500ML | Sigma |
| 2-phenyethanol (2pe) | $C_{8}H_{10}O$ | 77861-250ML | Sigma |
| isoamyl acetate (iso) | $C_{7}H_{14}O_{2}$ | W205508-SAMPLE-K | Sigma |

**Supplemental Table 2 – Antibodies used in this study**

**Primary antibodies**

| Antigen | Species | Dilution | Catalog No. | Supplier |
|---|---|---|---|---|
| SOX2 | Rat | 1/100 | 14-9811-82 | Invitrogen |
| CK14 | Rabbit | 1/100 | ab119695 | Abcam |
| OMP | Rabbit | 1/500 | ab183947 | Abcam |
| S100β | Rabbit | 1/500 | ab52642 | Abcam |
| DCX | Rabbit | 1/250 | ab18723 | Abcam |
| CRE | Guinea pig | 1/100 | 257-004 | Synaptic System |
| GFP | Goat | 1/500 | 601-101-215 | Rockland |

**Secondary antibodies**

Respective secondary antibodies raised against rat, rabbit, guinea pig and goat coupled with Alexa Fluor® or DyLight® fluorophores of excitation wavelengths 405nm, 488nm, 594nm, and 649nm were purchased from Jackson Immunoresearch. All secondary antibodies were used at 1/500 dilution.
